# Proteomic analysis of the role of the quality control protease LONP1 in mitochondrial protein aggregation

**DOI:** 10.1101/2021.04.12.439502

**Authors:** Karen Pollecker, Marc Sylvester, Wolfgang Voos

## Abstract

The mitochondrial matrix protease LONP1 is an essential part of the organellar protein quality control system. LONP1 has been shown to be involved in respiration control and apoptosis. Furthermore, a reduction in LONP1 level correlates with ageing processes. Up to now, the effects of a LONP1 defect were mostly studied by utilizing transient, siRNA-mediated knockdown approaches. We generated a new cellular model system for studying the impact of LONP1 on mitochondrial protein homeostasis by a CRISPR/Cas-mediated genetic knockdown (gKD). These cells show a stable reduction of LONP1 along with a mild phenotype characterized by absent morphological differences and only small negative effects on mitochondrial functions under normal culture conditions. To assess the consequences of a permanent LONP1 depletion on the mitochondrial proteome, we analyzed the alterations of protein levels by quantitative mass spectrometry, demonstrating small adaptive changes, in particular concerning mitochondrial protein biogenesis. In an additional proteomic analysis, we determined the temperature-dependent aggregation behavior of mitochondrial proteins and its dependence on a reduction of LONP1 activity, demonstrating the important role of the protease for mitochondrial protein homeostasis in mammalian cells. We identified a significant number of mitochondrial proteins that are affected by LONP1 activity concerning their stress-induced solubility. Taken together, our results suggest a very good applicability of the LONP1 gKD cell line as a model system for human ageing processes.

## Introduction

A balanced relationship between protein synthesis and degradation, called protein homeostasis or proteostasis in short, is essential for the fitness and survival of cellular systems. An important functional aspect of proteostasis is the specific degradation of damaged or aggregated proteins that are generated under stress conditions (1). While ageing, proteostasis systems lose their capacity and thus the risk of damage caused by aggregated proteins is increased (2). Accumulation of these aggregated proteins may further disrupt the proteostasis efficiency and can eventually lead to neurodegenerative diseases, like Alzheimer’s and Parkinson’s disease (3). As mitochondria have a central role in cellular metabolism, mitochondrial dysfunction is observed during ageing processes and also in many neuronal diseases (4,5). An essential question is whether disruption of mitochondrial protein homeostasis by aggregation is the cause of mitochondrial dysfunction under such pathological conditions. Increased ambient temperatures are a typical stressor that lead to an accumulation of damaged polypeptides in cells and also inside mitochondria (6). Elevated temperatures are widely used to study protein stability in model systems as they cause protein unfolding and aggregation, accompanied by decreased metabolic functions (6,7). Due to their endosymbiotic origin, mitochondria possess their own protein quality control system (mtPQC), which consists of dedicated proteases, chaperones and co-chaperones (8). This control system becomes upregulated by the so-called mitochondrial unfolded protein response (UPR^mt^) pathway (9). This mtPQC system is essential for maintenance of proteostasis in mitochondria, first by trying to refold misfolded polypeptides or, if this is not possible, by degradation of these damaged proteins. Typically, the degradation of misfolded proteins is performed by ATP-dependent proteases of the AAA+ (ATPases associated with a wide variety of cellular functions) family (8). Mitochondria contain four different AAA+ proteases, LONP1, CLPP (mitochondrial ATP-dependent CLP protease proteolytic subunit), AFG32/SPG7 (AFG-like protein 2/Paraplegin) and YME1L1 (ATP-dependent zinc metalloprotease) (8). One of the main proteases located in the mitochondrial matrix is the highly conserved protease LONP1(10–12), named also La (13) in bacteria as well as Pim1 in yeast (14). Like other AAA+-type serine-proteases, LONP1 forms a chambered protease complex for general protein degradation (15). Chambered proteases typically consist of an ATPase domain, mainly responsible for substrate recognition and unfolding, and a proteolytic domain where the active site is shielded from the environment. In case of LONP1 both domains are on the same polypeptide chain, while in other cases two separate proteins cooperate closely and form the active protease complex (16). In mitochondria, the Lon-type proteases are thought to closely cooperate with chaperones from the heat shock protein (HSP) 70 family to form a cooperative mtPQC network dealing with polypeptide misfolding situations (17). In contrast to the ubiquitin-proteasome system, a dedicated targeting system for substrate polypeptide degradation by LONP1 is lacking in mitochondria. Substrate proteins rather seem to be recognized due to an at least partially unfolded state (18,19), correlating with its protective role in the mtPQC system. Although a few genuine LONP1 substrate proteins, in particular under oxidative stress situations (20,21), have been identified so far. However, the full endogenous substrate spectrum of LONP1, in particular in mammalian cells, has not been fully established.

LONP1 has been shown to be up-regulated in response to external stressors, e.g. heat stress, serum starvation and oxidative stress situations (10,22). Moreover, the LONP1 level was found to be reduced in ageing cells, correlated with an accumulation of oxidatively damaged and aggregated proteins (21), as well as an irregular mitochondrial morphology (23). In yeast, a deletion of the *PIM1* gene was viable but resulted in a major defect in mitochondrial respiratory rates (14) and accumulation of oxidized polypeptides (24). In contrast, a complete LONP1 knock out resulted in embryonic lethality in mice at day 9.5 (25), showing that LONP1 is essential for the survival of mammalian organisms. A siRNA-mediated reduction of LONP1 level in cultured cells lead to an increase of oxidatively modified proteins, correlated with an appearance of electron-dense structures in the mitochondrial matrix, potentially representing aggregated polypeptides (23,26). Although these results establish a general role of the LONP1 protease in the protection of mitochondrial functional integrity, in particular in human cells, the details of its functions and the consequences of more subtle LONP1 defects on protein homeostasis in the mitochondrial matrix remain to be clarified.

Here we present the developments of a CRISPR/Cas-mediated genetic LONP1 knock down (gKD) in HeLa cells that is based on small deletions in the promoter sequence of the gene. A gKD approach was preferred over a full knock out as a stable reduction of cellular LONP1 amounts might simulate different pathological situations that are connected to a decrease of mitochondrial protein homeostasis. For an overview of the cellular effects of an overall reduction of LONP1 amounts, general mitochondrial functions were analyzed. In order to obtain a comprehensive picture of the role of LONP1 on mitochondrial protein homeostasis as our main focus, we analyzed the changes in the mitochondrial proteome resulting from a reduction of the protease in LONP1 gKD cells via quantitative mass spectrometry (qMS) of isolated mitochondria. In a second qMS approach, we directly characterized the temperature-dependent aggregation behavior of mitochondrial proteins in wild type (WT) and the LONP1-depleted cellular background and identified novel mitochondrial protein targets depending on the activity of LONP1.

## Experimental Procedures

### CRISPR/Cas mediated LONP1 knock down and cell culture conditions

The LONP1 gKD was achieved by a CRISPR/Cas9 mediated genome alteration (27) targeting a region shortly upstream of the LONP1 start codon in HeLa cells. The selected guide sequence (ACGTGCGTTTCGCGGCGAG) was cloned into the CRISPR/Cas9 containing vector PX459 V2.0. pSpCas9(BB)-2A-Puro (PX459) V2.0 was a gift from Feng Zhang (Addgene plasmid #62988; http://n2t.net/addgene:62988; RRID:Addgene_62988). HeLa WT cells were transfected with 1 μg/ml plasmid using TurboFect transfection reagent (Thermo Fisher) according to manufacturer’s instruction and selected with 2 μg/ml puromycin for four days. Transfected cells were diluted to a single cell colony and cultured. The procedure led to a Cas9-induced DNA double strand break 8 nt upstream of the LONP1 start codon. The successful deletion was confirmed by MiSeq-sequencing of selected LONP1 gKD cell clones, performed by the Next Generation Sequencing Core Facility (University Hospital Bonn).

HeLa WT and HeLa LONP1 gKD cells were cultured in Roswell Park Memorial Institute 1640 medium (RPMI medium, Invitrogen), containing 10% (v/v) fetal calf serum (FCS, Invitrogen), 2 mM L-glutamine, 100 units/ml penicillin and 100 μg/ml streptomycin at 37°C in 5% CO_2_. Cells were grown on tissue culture dishes and passaged at ratio 1:5 or 1:10 by trypsin treatment every 72 to 96 h.

### Western blot analysis

For Western blot analysis, mitochondria were isolated as described previously (28). Whole cells were lysed with lysis buffer (0.5% Triton-X100 (v/v); 30 mM Tris pH 7.4; 5 mM EDTA; 200 mM NaCl; 0.5 mM PMSF; Protease Inhibitor cocktail, Roth). 15 μg total protein from cell lysates or isolated mitochondria were dissolved in Laemmli buffer (2% (w/v) SDS; 10% (v/v) glycerol, 60 mM Tris pH 6.8; 5% (w/v) β-mercaptoethanol, 0.02% (w/v) bromphenol blue) were loaded on a 15% SDS-PAGE and electrically transferred via Western blot on a PVDF-membrane. Afterwards, the membrane was blocked with 5% milk in TBS-Tween (1.25 M NaCl; 200 mM Tris/HCl, pH 7.5; 0.1% (v/v) Tween-20) or Dekosalt-Tween (15 mM NaCl; 5 mM Tris; 0.1% (v/v) Tween-20), and decorated with different antibodies αLONP1, αTRAP1, αCLPX (own work; unpublished); αSDHA (ProteinTech 14865-1-AP); αTIMM23 (BD Bioscience 611222); αmtHSP70 (Stressgen SPS-825); αCLPP (ProteinTech 15698-1-AP); αHSP60 (Santa Cruz SC-13966); αGAPDH (AK-online ABIN 274251). After incubation over night at 4°C, the membrane was decorated with the appropriate peroxidase-coupled secondary antibodies (Sigma-Aldrich). The chemiluminescence reaction with respective substrate solution (Serva) was detected with a camera system (LAS-4000, FUJI).

### Measurement of membrane potential and ATP content in isolated mitochondria

Membrane potential was measured with 1 μM tetramethylrhodamine (TMRE, excitation: 540 nm, emission: 595 nm; Sigma Aldrich) as described before (6), here 40 μg of freshly isolated mitochondria were used. For negative control, mitochondria were treated with 1 μM valinomycin. TMRE fluorescence was detected in a microplate reader (Infinite M200 Pro, Tecan). In order to determine the ATP content, a luciferase-based assay (ATP Determination kit, Invitrogen) was used. 75 μg mitochondria were lysed with 0.5% triton X-100, according manufacture’s protocol. The luciferase activity was measured in a microplate reader (Infinite M200 Pro, Tecan).

### Flow cytometry analysis of reactive oxidative species

To determine the generation of reactive oxidative species (ROS) in intact cells, the mitochondria-specific superoxide indicator MitoSOX (Thermo Fischer Scientific) was used as described previously (6). For positive control cells were stressed with 0.1 mM menadione for 60 min at 37°C and 5% CO_2_. Afterwards, treated and untreated cells were incubated for 20 min with 5 mM MitoSOX reagent at 37°C and 5% CO_2_ at ambient O_2_ concentrations. Cells were harvested after trypsinization and washed thrice with cold PBS containing 0.2% (w/v) BSA. Fluorescence intensities of 50,000 cells for each sample were measured by using a flow cytometer (CyFlow space CY-S3001; Partec).

### Immunofluorescence microscopy for mitochondrial morphology

Cells were seeded on sterile cover slides and allowed to get attached overnight. After 5 min of fixation with 4% formaldehyde and 10% sucrose solution in PBS at 37°C, the cell membrane was permeabilized with a 0.5% (v/v) Triton X-100 in PBS for 12 min and blocked with 2% BSA in PBS for 1 h at room temperature. Afterwards, cells were decorated with αSDHA (ProteinTech, 14865-1-AP) and incubate at 4°C overnight. After washing with PBS, cells were decorated with an fluorescent αrabbit-antibody (αrabbit Cy3, M30010, Invitrogen). Cover slides were mounted with mounting medium containing DAPI (Roth), after washing twice with PBS and once with H_2_O. For microscopy, the EVOS_FI_ Cell Imaging System (AMG, Germany) was used.

### *In organello* translation of mitochondrial encoding proteins

The translation of mitochondria-encoded proteins in intact isolated mitochondria was performed essentially as described (6). 50 μg fresh isolated mitochondria were resuspended in translation buffer (645 mM sorbitol, 160 mM KCl, 16 mM KPi pH 7.2, 21.5 mM Tris pH 7.4, 13.5 mM MgSO_4_, 3.3 μg/μl BSA, 21.5 mM ADP, 0.53 mM GTP, 13.8 mM creatine phosphate, 4 μg/ml creatine kinase, 1.2 mg/ml α-ketoglutarate, 14 μM amino acid mix w/o methionine (Promega)). As negative control, mitochondria were preincubated with chloramphenicol (f. c. 100 μg/ml) or cycloheximide (f. c. 100 μg/ml) for 3 min at 30°C. To label newly synthesized mitochondrial proteins, [^35^S]-methionine/cysteine (f. c. 22 μCi/μl) was added. After incubating for 45 min at 30°C, labelling was stopped by adding MOPS-Met buffer (1 M MOPS pH 7.2, 200 mM methionine, f. c. 50 mM). The mitochondria were incubated again for 30 min at 30°C and washed with washing buffer (0.6 M sorbitol, 1 mM EDTA, 5 mM methionine). The mitochondrial pellet was resuspended in Laemmli buffer, then applied on a 15% SDS-PAGE containing 1.1 M urea. Afterwards, the newly synthesized proteins were analyzed by Western blot and autoradiography.

### Mitochondrial preprotein import *in vivo*

A potential mitochondrial import defect in intact cells was assessed by the appearance of unprocessed preprotein signals. In control cells, import reactions were blocked *in vivo* by disrupting the inner membrane potential (Δψ). Cells were treated with the uncoupling chemicals 1 μM valinomycin for 4 h under standard growth condition. Then cells were lysed in 0.5% (v/v) Triton X-100 lysis buffer (30 mM Tris pH 7.4, 200 mM NaCl, 5 mM EDTA pH 8, 0.5 mM PMSF and 1x protease inhibitor cocktail (Roth). 20 μg of protein lysate were taken up in Laemmli buffer and loaded onto a 10% SDS-PAGE and analyzed by Western blot and immunodecoration with αTRAP1, αLONP1 (own work) and αGAPDH (AK-online ABIN 274251).

### *In organello* degradation of imported, radioactive labelled reporter proteins

Mitochondria of WT and LONP1 gKD cells were isolated and radioactive labelled preproteins were imported as described previously (28). Two different preproteins were used: TRAP1 and MDH2, which were each translated and transcribed *in vitro* with TNT-coupled reticulocyte lysate system (Promega) and imported for 10 min at 30°C in import buffer. Import was stopped with 0.5 μM valinomycin treatment. Afterwards, 25 μg mitochondria per sample were resuspended in degradation buffer with an ATP-regenerating system (0.25 M sucrose, 20 mM HEPES pH 7.6, 80 mM KAc, 5 mM MgAc, 5 mM glutamate, 5 mM malate, 1 mM DTT, 5 mM KPi, 2 mM ATP, 10 mM creatine phosphate, 75 μg/ml creatine kinase, 0.02 μg/ml trypsin inhibitor). 1 mM menadione (dissolved in ethanol) was added to generate ROS-stressed mitochondria, in control samples, the same amount of ethanol was added. Degradation samples were incubated for up to 480 min at 30°C. All samples were precipitated with TCA (f. c. 14.4% (w/v)), and pellets were resuspended in Laemmli buffer. The reporter proteins were detected after SDS-PAGE and Western blot through autoradiography. Smaller degradation fragments appearing over time were quantified with Multi Gauge software (FUJI). To determine the degradation rate, the amounts of the degradation fragments were plotted against the incubation time. The rate was calculated by using the slope of the regression line and and was normalized to the untreated WT sample.

### Mitochondrial aggregation assay

For analysing heat-induced aggregation of mitochondrial proteins, mitochondria were isolated from cultured HeLa cells as described before (28). 30 μg mitochondria were taken up in resuspension buffer (250 mM sucrose; 20 mM HEPES, pH 7.6; 80 mM KAc; 5 mM MgAc; 5 mM glutamate; 5 mM malate; 1 mM DTT) and heat stressed for 20 min at 25°C, 37°C, 42°C or 45°C. After lysis with Triton-lysis buffer (0.5% Triton X-100 (v/v); 30 mM Tris pH 7.4; 5 mM EDTA; 200 mM NaCl; 0.5 mM PMSF; Protease Inhibitor cocktail, ROTH), mitochondrial lysates were centrifuged at 125,000xg for 45 minutes to separate supernatant (soluble proteins) and pellet with aggregated proteins (6,28). Proteins in supernatant were precipitated with TCA (f.c. 14.4% (w/v)) and both pellets were resuspended in Laemmli buffer. Samples were loaded onto an 12.5% SDS-PAGE and analyzed by Western blot and immunodecoration with αACO2 (Sigma Aldrich HP001097), αLONP1 (own work), αHSP60 (Santa Cruz 13966), αTUFM Protein (Tech 11701-1-AP), αTSFM (Sigma Aldrich HPA024087), αTFAM (NEB 80760S) and αTIMM23 (BD. Bioscience 80760).

### SILAC labelling and mass spectrometry analysis

Cells were grown in labelling medium containing 0.1 mg/ml normal lysine or 0.1 mg/ml ^13^C_6_-lysine (heavy) lysine (Gibco) for 4 passages (~2 weeks). Afterwards, mitochondria were isolated as described before. 30 μg WT and 30 μg LONP1 gKD mitochondria were mixed and heat stressed at 42°C or incubated at 25°C for 20 min. Sedimentation of aggregated protein were described in mitochondrial aggregation assay. Together with non-treated mitochondria, samples were loaded on a 12% SDS-PAGE. Afterwards, proteins were cut out of the acrylamide gel and analyzed by mass spectrometry as follows.

#### Peptide preparation

Gel slices were subjected to tryptic in gel digestion (29). In brief, slices were washed consecutively with water, 50% acetonitrile (ACN), and 100% ACN. Proteins were reduced with 20 mM DTT in 50 mM ammonium bicarbonate and alkylated with 40 mM acrylamide (in 50 mM bicarbonate). The slices were washed again and dehydrated with ACN. Dried slices were incubated with 330 ng sequencing grade trypsin at 37°C overnight. The peptide extract was separated and remaining peptides extracted with 50% ACN. Peptides were dried in a vacuum concentrator and stored at −20°C.

#### LC-MS measurements of peptides

Peptides were dissolved in 10 μl 0.1% formic acid (FA) and 3 μl were injected onto a C18 trap column (20 mm length, 100 μm inner diameter, ReproSil-Pur 120 C18-AQ, 5 μm, Dr. Maisch GmbH, Ammerbuch-Entringen, Germany) made inhouse. Bound peptides were eluted onto a C18 analytical column (200 mm length, 75 μm inner diameter, ReproSil-Pur 120 C18-AQ, 1.9 μm, with 0.1% formic acid as solvent A). Peptides were separated during a linear gradient from 5% to 35% solvent B (90% acetonitrile, 0.1% FA) within 90 min at 300 nl/min. The nano HPLC was coupled online to an LTQ Orbitrap Velos mass spectrometer (Thermo Fisher Scientific, Bremen, Germany). Peptide ions between 330 and 1600 m/z were scanned in the Orbitrap detector with a resolution of 60,000 (maximum fill time 400 ms, AGC target 10^6^). The 22 most intense precursor ions (threshold intensity 3000, isolation width 1.0 Da) were subjected to collision induced dissociation (CID, normalized energy 35) and analyzed in the linear ion trap. Fragmented peptide ions were excluded from repeat analysis for 15 s.

#### Data analysis

For peptide analysis, raw data processing and database searches were performed with Proteome Discoverer software 2.3.0.523 (Thermo Fisher Scientific). Peptide identifications were done with an in-house Mascot server version 2.6.1 (Matrix Science Ltd, London, UK). MS2 data were searched against human sequences in SwissProt (release 2017_10). Precursor Ion m/z tolerance was 9 ppm, fragment ion tolerance 0.5 Da (CID). Tryptic peptides with up to two missed cleavages were searched. Propionamide on cysteines was set as a static modification. Oxidation of methionine and ^13^C_6_ label on lysine were allowed as dynamic modifications. Mascot results were assigned q-values by the percolator algorithm (30) version 3.02.1 as implemented in Proteome Discoverer. Spectra with identifications below 1% q-value were sent to a second round of database search with semitryptic enzyme specificity (one missed cleavage allowed) and 10 ppm MS1 mass tolerance (propionamide dynamic on Cys). Proteins were included if at least two peptides were identified with <1% FDR. Only unique peptides were included in protein quantification.

### Statistical analysis

All statistical analyses were performed with GraphPad prism 7 and show mean values and standard error of the mean (SEM). Statistical significance was determined by Students T-test of at least 3 independent replicates.

## Results

### General effects of a reduced LONP1 level

We created stable genetic knock down (gKD) of the mitochondrial matrix protease LONP1 by a CRISPR/Cas-mediated process in cultured HeLa cells (Fig. S1A). Genomic sequencing showed a 7 nt deletion upstream close to *LONP1* start codon (Fig. S1B). The Western blot analysis of cellular protein levels from LONP1 gKD cells showed a strong reduction of LONP1 amounts compared to WT cells both in total cell lysates (Fig. 1A, top panel) as well as in isolated mitochondria (Fig. 1B, top panel, Fig. S1C). The subsequent quantitative MS characterization of LONP1 gKD mitochondria (see below) confirmed a reduction of the protease to about 25% of the WT levels (Fig. 1C). Furthermore, we determined the amounts of typical control proteins from different mitochondrial localizations to get first impression of the general effect of a LONP1 depletion on the mitochondrial protein composition (Fig. 1A, B). The mitochondrial outer membrane protein, TOMM70 (mitochondrial import receptor subunit TOMM70) was slightly increased in total cell lysate as well as in isolated mitochondria. TIMM23 (mitochondrial import inner membrane translocase subunit TIMM23), a protein located in mitochondrial inner membrane and providing the pore for preprotein translocation, exhibited a similar behavior. As LONP1 functions in mtPQC, we determined the amounts of the most prominent chaperone proteins from the mitochondrial matrix, HSP60 (heat shock protein 60 kDa) and mtHSP70 (mitochondrial heat shock protein 70 kDa), which were not altered, indicating the absence of a UPR^mt^ as a response to the LONP1 depletion. However, amounts of the alternative protease component in the mitochondrial matrix, CLPP, were strongly reduced after gKD of LONP1, while its partner chaperone protein, CLPX (mitochondrial ATP-dependent CLP protease ATP-binding subunit, CLPX-like), was not changed. Amounts of the reactive oxidative species (ROS)-protective enzyme manganese superoxide dismutase (SOD2) located in the mitochondrial matrix were slightly reduced in LONP1 gKD mitochondria. Due to the connection of LONP1 and mtDNA expression, we tested possible consequences of LONP1 depletion on oxidative phosphorylation by measuring the ATP content of isolated mitochondria with a luciferase-based assay. Indeed, ATP concentrations were reduced to about 40% (Fig. 1D) in LONP1 gKD mitochondria. The mitochondrial inner membrane potential, generated by the respiratory chain, was measured by using the mitochondria-specific, fluorescent dye TMRE. Intact WT and LONP1 gKD cells were incubated with TMRE and the fluorescence of the cells was assayed by flow cytometry. As negative control, we diminished the membrane potential by treating cells with 0.1 μM valinomycin for 10 min prior to TMRE staining. Isolated LONP1 gKD mitochondria showed a similar membrane potential as WT mitochondria (Fig. 1E). An accumulation of ROS in the mitochondria has been often observed under proteostatic stress conditions. However, mitochondrial ROS levels, monitored by the mitochondria-specific superoxide reactive dye MitoSOX, as such were not increased in our LONP1 gKD cells. In contrast, after a short treatment of the cells with the ROS-generating chemical menadione, the gKD cells exhibited higher ROS levels (Fig. 1F), most likely a direct consequence of the reduced protein level of SOD2 we detected in the Western blot analysis. To test for morphological changes, cells were grown on cover slips, fixed and decorated with an antibody against the inner membrane respiratory chain complex II component SDHA (succinate dehydrogenase subunit A, flavoprotein) to visualize the mitochondrial tubular network. Both WT and LONP1 gKD cells showed a comparable normal tubular mitochondrial structure and no stress-dependent fragmentation of mitochondria was observed in LONP1 gKD (Fig. 1G). Taken together, the LONP1 gKD cells are morphologically unremarkable and showed only a mild functional phenotype.

**Figure 1.**
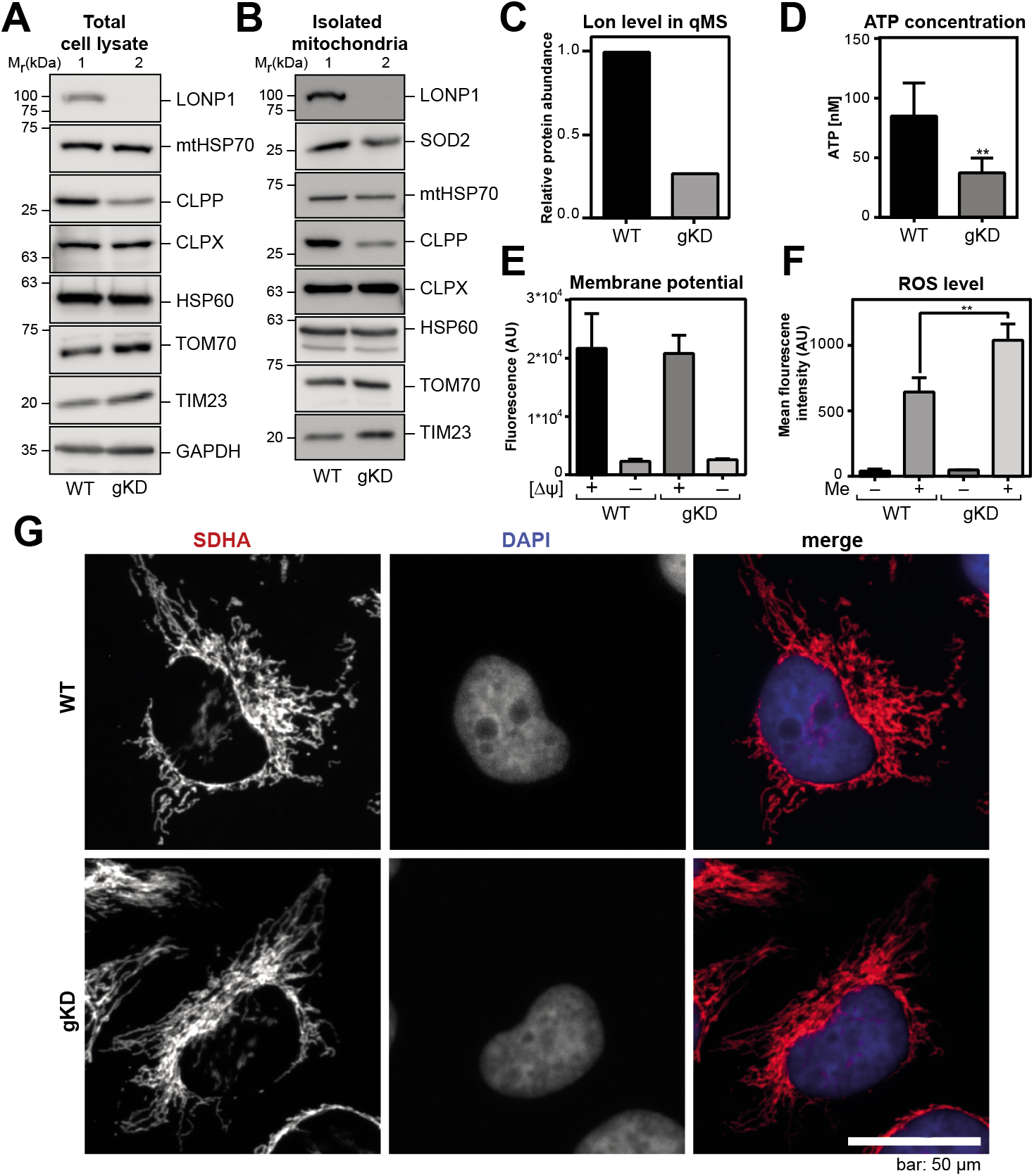
Characterization of newly designed LONP1 gKD cells. Detection of different mitochondrial and cytosolic proteins in total cell lysates (A) and isolated mitochondria (B) of WT and LONP1 gKD cells by SDS-PAGE, Western blot and immunodetection. Antibodies against the indicated proteins were used as listed in Material and Methods. Indicated are the corresponding relative molar masses of the respective SDS-PAGE marker proteins. GAPDH and HSP60 signals were used as loading control. (C) LONP1 protein levels (given as ratio to WT) in isolated mitochondria of WT (black) and LONP1 gKD (gray) cells used for quantitative MS analysis after two weeks labelling with normal or heavy isotope-labelled arginine. (D) Determination of ATP concentration in isolated and lysed WT (black) or LONP1 gKD (gray) mitochondria. ATP concentration was determined by using a luciferase-based assay as described. Shown are mean values ± SEM of three independent experiments (n=3), ** p<0.01. (E) Detection of mitochondrial inner membrane potential (Δψ) in isolated mitochondria of WT (black) and LONP1 gKD (dark gray) cells, using the potential-dependent fluorescent dye TMRE. As control, mitochondria were pre-treated with 1 μM valinomycin for 10 min at room temperature. Shown are mean values ± SEM (n=3) of TMRE fluorescence emission intensities. (F) Measurement of mitochondrial ROS amounts in WT (black) and LONP1 gKD (dark gray) cells by flow cytometry analysis using the mitochondria-specific superoxide dye MitoSOX. As positive control, cells were pre-treated with 0.1 mM menadione (Me) for 60 min (WT, gray; LONP1 gKD, bright gray). Shown are mean values ± SEM (n=3) of MitoSOX fluorescence emission intensity; ** p<0.01. (G) Immunofluorescence analysis of mitochondrial morphology by using fluorescence microscopy of fixed WT and LONP1 gKD cells. Anti-SDHA antibodies were used to visualize the mitochondrial network (red), nuclei were stained by DAPI (blue). Upper panel, HeLa WT cells; lower panel, LONP1 gKD cells. Bar: 50 μm.

Mitochondrial transcription and translation were analyzed by radioactive pulse-chase labelling of mitochondria-encoded proteins. Isolated mitochondria were allowed to synthetize proteins in presence of ^35^S-methionine/cysteine for 45 min at 30°C. Afterwards, radioactive mitochondrial proteins were detected through SDS-PAGE and autoradiography. One of 13 mitochondrial encoded proteins, ND4 (NADH-ubiquinone oxidoreductase chain 4) exhibited a slightly more efficient translation compared to WT. In contrast, translation rate of ATP8 (ATP synthase protein 8) level was strongly reduced in LONP1 gKD mitochondria in comparison to WT (Fig. 2A; lanes 1 and 4). As control, a preincubation with the translational inhibitors chloramphenicol (Fig. 2A; lanes 2 and 5) and cycloheximide (Fig. 2A; lanes 3 and 6) resulted in a strongly reduced labeling reaction, indicating the specificity of the translation reaction to mitochondria.

**Figure 2.**
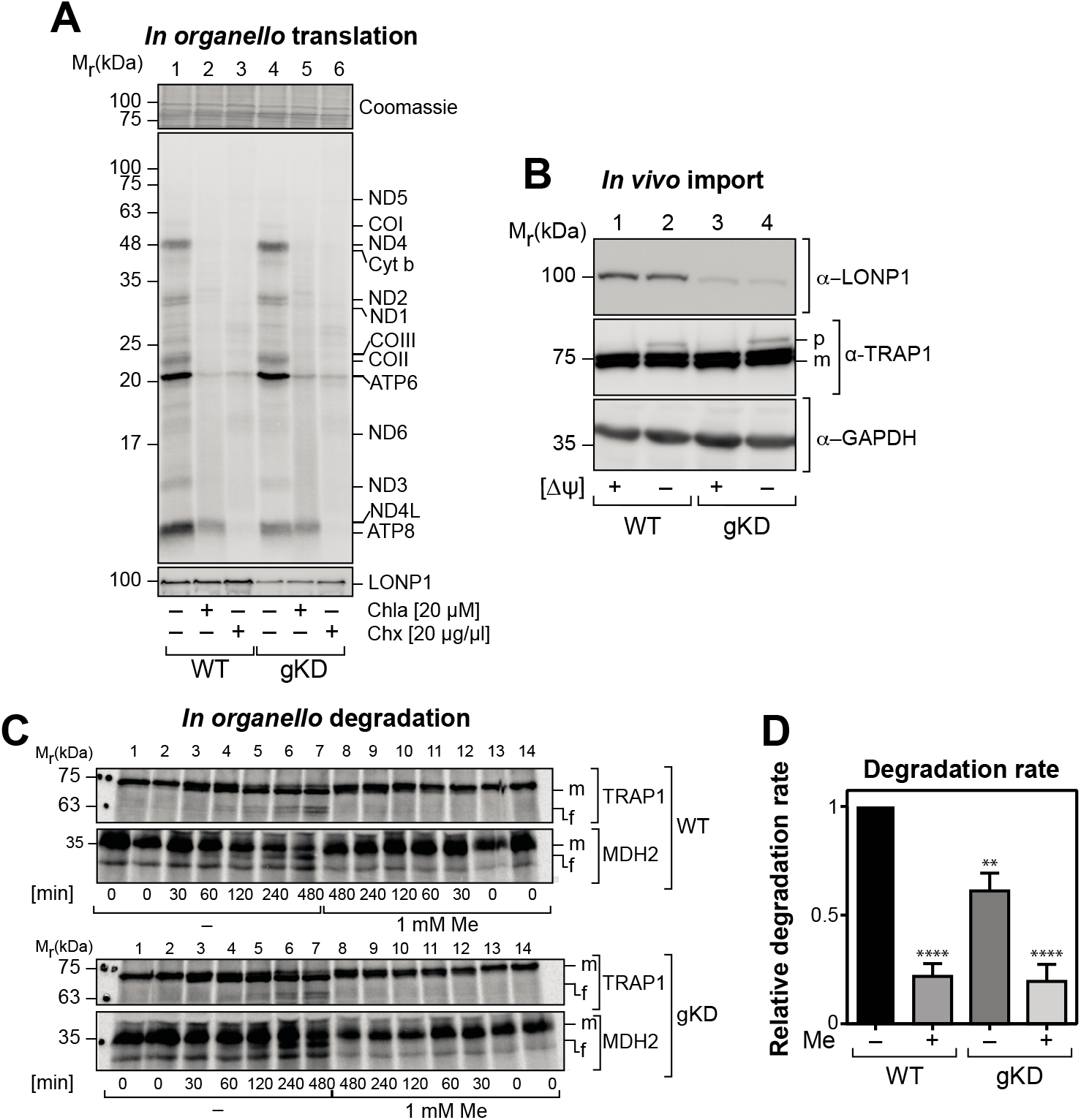
Mitochondrial protein biogenesis and degradation in LONP1 gKD mitochondria. (A) Detection of translation in isolated mitochondria of WT and LONP1 gKD mitochondria after incubation for 45 min with [35S]-methionine and cysteine at 30°C. Newly synthesized mitochondrial proteins were detected by 15% Urea-SDS-PAGE, Western blot and autoradiography (middle panel). Upper panel shows section of commassie-stained membrane as loading control and lower panel shows LONP1 amounts detected by immunodecoration. As translation control, mitochondria were pre-treated with the translation inhibitors Cycloheximid (CHX; 20 μg/μl) and Chloramphenicol (Chla; 20 μM). Indicated are the corresponding relative molar masses of the respective SDS-PAGE marker proteins. (B) Analysis of mitochondrial import in intact cells assessed by an accumulation of endogenous preproteins. As control, import was blocked by 1 μM valinomycin treatment for 60 min (-Δψ, lane 2 and 4). Un-processed precursor (p) and imported mature (m) forms of the mitochondrial matrix protein TRAP1 were detected with SDS-PAGE, Western blot and immunodetection. As controls, the mitochondrial protein LONP1 and the cytosolic protein GAPDH are shown. Antibodies against the indicated proteins were used as listed in Material and Methods. Indicated are the corresponding relative molar masses of the respective SDS-PAGE marker proteins. (C) Detection of in organello degradation of imported, radioactively labelled preproteins TRAP1 and Mdh2. Import into isolated WT (upper panel) and LONP1 gKD (lower panel) mitochondria was stopped after 10 min and degradation rate was analyzed over a further incubation period of 480 min. For ROS stress conditions, mitochondria were incubated with 1 mM menadione (Me) during the degradation incubation. Imported and processed proteins (m) as well as degradation fragments (f) were detected by SDS-PAGE and autoradiography. Shown is a representative experiment. (D) Quantitative assessment of degradation rates as obtained in (C). The rate of the increase in band intensities of the degradation fragment (f) over time was determined and normalized to WT (untreated). Shown are mean values ± SEM (n=3) of degradation rate relative to WT (black), Significance values are calculated relative to WT control; **** p<0.0001; ** p<0.01.

We also assessed the impact of LONP1 gKD on protein import as a main feature of mitochondrial protein biogenesis. Here, we analyzed the import of endogenous mitochondrial protein TRAP1 (tumor necrosis factor type 1 receptor-associated protein) in intact cells by the detection of the full-length non-processed form and the fully imported processed mature variant. As control, cells were treated with valinomycin treatment for 4 h, depleting the inner membrane potential. During this time, the precursor form TRAP1 accumulated in cytosol and became visible as a higher molecular-weight signal (Fig. 2B, lanes 2 and 4). With full membrane potential, we could not detect any accumulation of precursor forms neither in WT nor in LONP1 gKD cells (Fig. 2B, lanes 1 and 3). Loading controls were represented by the cytosolic protein GAPDH and LONP1 itself. Thus, no major preprotein import defect due to the reduction of the LONP1 protein amounts was observable. As the main proof-reading protease in the mitochondrial matrix, a LONP1 reduction may affect the degradation of imported polypeptides. For this purpose, the radiolabeled precursor proteins TRAP1 and MDH2 (malate dehydrogenase 2) were imported as reporter proteins *in organello* for 10 min into mitochondria isolated from WT and LONP1 gKD cells. After stopping the import reaction by an addition of valinomycin, the degradation of the imported polypeptides in the mitochondrial matrix was followed over a period of 480 min by detecting radioactive degradation fragments using SDS-PAGE and autoradiography (Fig. 2C). Degradation rates were indirectly calculated by quantifying the increase of the newly appearing degradation fragment (Fig. 2C; f, fragment). After 240 min, a difference in the degradation rate of TRAP1 between WT and LONP1 gKD mitochondria could be detected. At the end of the incubation the amount of the TRAP1 degradation fragment was twice as high as in LONP1 gKD mitochondria (Fig. 2C, upper panel). Similarly, after 480 min a comparable difference concerning the degradation of imported MDH2 was measured (Fig. 2C, lower panel). In addition, both WT and LONP1 gKD mitochondria after menadione treatment – simulating proteotoxic stress conditions - showed a very low degradation rate, with non-detectable differences between the two cell types. Quantification of degradation rates relative to WT showed a decrease in LONP1 gKD mitochondria by ~60% (p = 0.032; Fig. 2D). The reduction of LONP1 hence resulted in a significant decrease in proteolytic capacity in the mitochondrial matrix.

In principle, the LONP1 gKD cells are viable and exhibit slight adaptations to the loss of LONP1, also preserving normal mitochondrial morphology. The mitochondria are still functional in terms of mitochondrial protein import and membrane potential. However, there is a decrease in mitochondrial ATP production and in the degradation rate of imported reporter proteins. The slowed but not abolished rate of degradation indicates that the ~25% remaining LONP1 levels are still functional at least principally able degrade proteins in mitochondria. In addition, there is also a slight reduction in the translation of specific mitochondria-encoded polypeptides, indicating a connection of translation and LONP1 function.

### Proteome changes after LONP1 gKD

To analyze long-term effects of a LONP1 depletion, we analyzed the changes in the mitochondrial proteome by quantitative MS (qMS) measurements of mitochondria isolated from WT and LONP1 gKD cells utilizing a SILAC approach. Cells were labelled for two weeks with the non-radioactive nitrogen isotope-containing the amino acid arginine (heavy for LONP1 gKD; normal for WT). Subsequently, mitochondria were isolated by the standard procedure and total protein extracts were separated by SDS-PAGE. Proteins were extracted from the gel, digested with trypsin and analyzed by qMS to determine the abundance of all detected mitochondrial proteins in comparison between WT and LONP1 gKD mitochondria. As we used a crude mitochondrial preparation procedure in this qMS analysis, we detected a total number of 3302 individual protein hits. By comparison with the cellular localization information available on the protein database UniProtKB (https://www.uniprot.org/), about 20% of the detected polypeptides could be unequivocally assigned as mitochondrial proteins in both samples. All non-mitochondrial proteins were eliminated from further analysis. The change in abundance for each individual protein was calculated as the quotient of proteins abundance values from WT and LONP1 gKD.

In total, 205 mitochondrial proteins were slightly but significantly increased in the LONP1 gKD mitochondria (up to 2.5 times, Fig. 3A, grey circles) and 15 were found to be increased up to ten times (Fig. 3A, dark grey circles). 12 proteins were strongly increased more than ten times (Fig. 3A, black circles). MRPL54 (mitochondrial 39S ribosomal protein subunit L54), MPC (mitochondrial pyruvate carrier) and BAX (apoptosis regulator BAX) were increased up to 20 times. MRPL53 (mitochondrial 39S ribosomal protein subunit L53), ATPD (mitochondrial ATP-synthase subunit delta), NDUFA4 (cytochrome *c* oxidase subunit) and ATP5MD (mitochondrial ATP synthase membrane subunit) were increased even stronger (Table S1). Notably, the amounts of previously published LONP1 substrates, like ACO2 (mitochondrial Aconitase 2) (21) and TFAM (transcription factor A, mitochondrial) (31), did not show significant differences in LONP1 gKD mitochondria.

**Figure 3.**
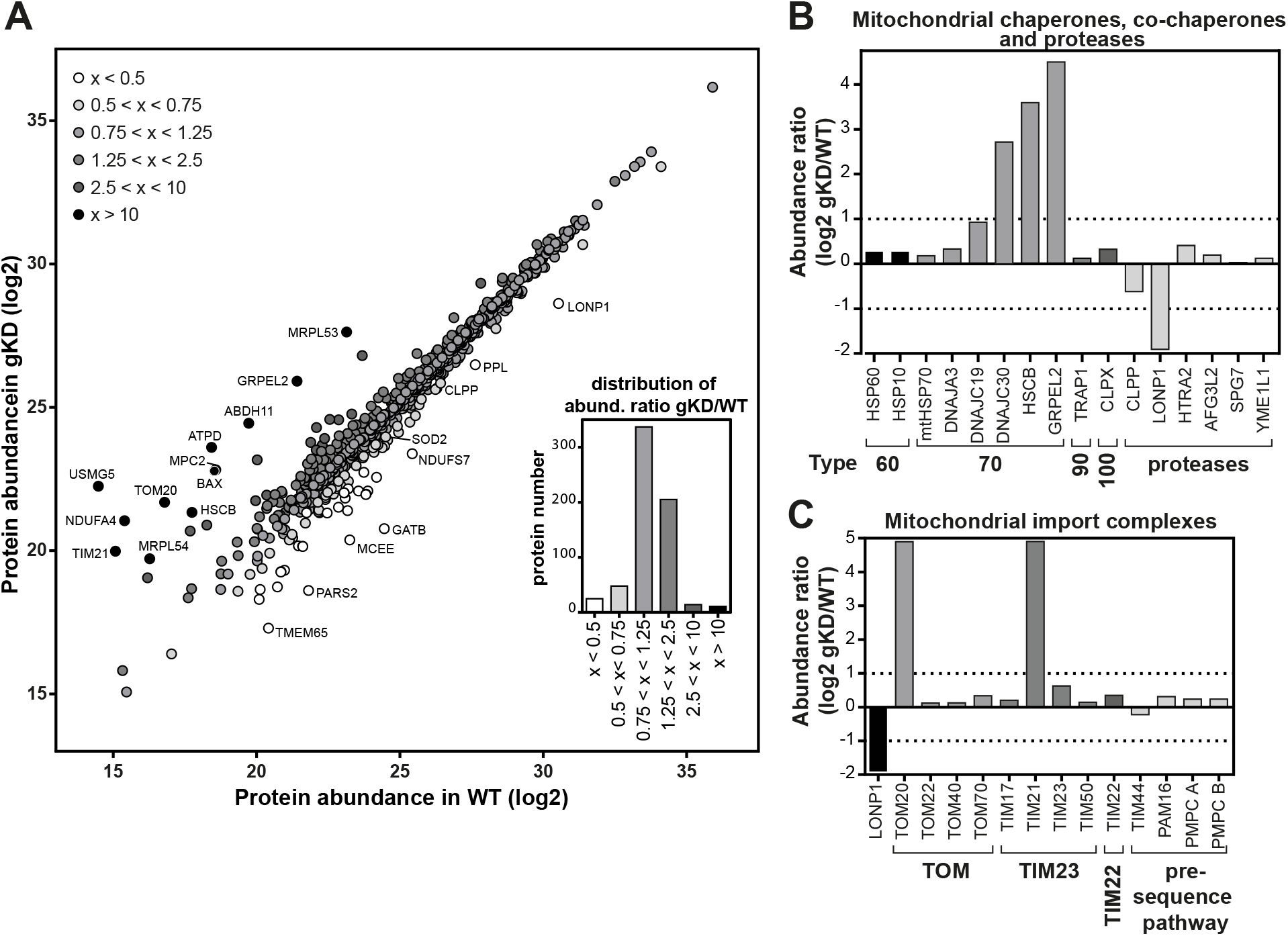
Changes in mitochondrial proteome after LONP1 gKD. (A) Total protein abundances (log2) of all detected mitochondrial proteins as obtained by qMS analysis of isolated WT and LONP1 gKD mitochondria. Inset shows numbers of proteins and their abundance ratios in gKD mitochondria relative to WT. Proteins strongly reduced in gKD (less than factor 0.5) are labeled white, light gray shows proteins with a factor between 0.5 and 0.75. Unchanged proteins (ratio between 0.75 and 1,25) are labelled gray, protein increased in gKD up to 10 times dark gray and over 10 times black. (B) Protein level differences (given as log2 ratio of gKD to WT) of selected members of the mitochondrial PQC system. Shown are the chaperones and co-chaperone of the indicated families and the major AAA+ proteases. (C) Protein level differences of selected members of the mitochondrial subunits of preprotein translocase systems of the outer membrane (TOM), inner membrane (TIM23 and TIM22 complex) and matrix-localized components of pre-sequence pathway compared to LONP1 (black).

Although the main chaperones in the mitochondrial matrix were essentially unaltered, three members of the mitochondrial HSP70 chaperone machinery family were strongly increased (Fig. 3B), HSCB (iron-sulfur cluster co-chaperone protein), GRPEL2 (mitochondrial GrpE protein homolog 2) and DNAJC19 (mitochondrial import inner membrane translocase subunit TIMM14). All three proteins operate as HSP70 co-chaperones (8) but with different functions. HSCB is involved in the iron-sulfur cluster assembly, while GRPEL2 and DNAJC19 are essential proteins associated with the preprotein import motor complex and required for the translocation of precursor proteins into the mitochondrial matrix. Indeed, two other components of the preprotein import translocase complexes showed a more than ten-fold increase, the preprotein receptor TOMM20 (about 30-fold), a subunit of TOM (translocase of the outer membrane) complex, and also TIMM21, a subunit of the TIMM23 (translocase of the inner membrane) complex (Fig. 3C).

Overall, the number of proteins with a lower amount in LONP1 gKD mitochondria was comparably small. We found 25 mitochondrial proteins, which were decreased by more than a factor of 0.5 in LONP1 gKD mitochondria (Fig. 3A, white circles). Two components of t-RNA synthesis, GATB (mitochondrial Glutamyl-tRNA aminotransferase subunit b) and PARS2 (a probable mitochondrial proline t-RNA ligase) were strongly reduced to less than 10% of WT protein level. 48 proteins were decreased to under 75% of WT level (Fig. 3A, light gray circles), here SOD2 - part of ROS stress response - was a prominent example. Interestingly, the other protease enzyme of the mitochondrial matrix, CLPP, was also found to be reduced to ~65% (Fig. 3B), confirming the Western blot analysis described above (Fig. 1A, B).

In summary, the majority of mitochondrial proteins showed only slight changes in abundance as a result of the permanent depletion of the LONP1 protease, correlating with the mild phenotype of LONP1 gKD cells. The observation that typical substrate proteins were not increased indicates that the LONP1-mediated protein degradation does not seem to play a prominent role under the used culture conditions. However, specific protein subgroups with a strong increase compared to WT mitochondria, in particular components of the mitochondrial HSP70 system and protein import machinery may indicate a potential compensation reaction to the LONP1 reduction.

### LONP1-dependent aggregation behavior

As a reduction of LONP1 activity should result in the accumulation of damaged polypeptides concomitant with their potential aggregation, we used a second proteomic approach to analyze the stress-dependent aggregation behavior of mitochondrial proteins. Again, cultures WT and LONP1 gKD cells were incubated with normal and heavy nitrogen isotope-containing arginine, respectively. After two weeks culture, mitochondria were isolated according to standard procedures, mixed, and subjected to a heat-stress treatment for 20 min at 42°C or at 25°C for control conditions. To separate soluble and aggregated proteins, mitochondria were lysed completely with a non-denaturing detergent under native conditions and centrifuged at 125,000xg (Fig. 4A) to recover potentially aggregated polypeptides. After a washing step, the sedimented proteins were analyzed by SDS-PAGE and qMS.

**Figure 4.**
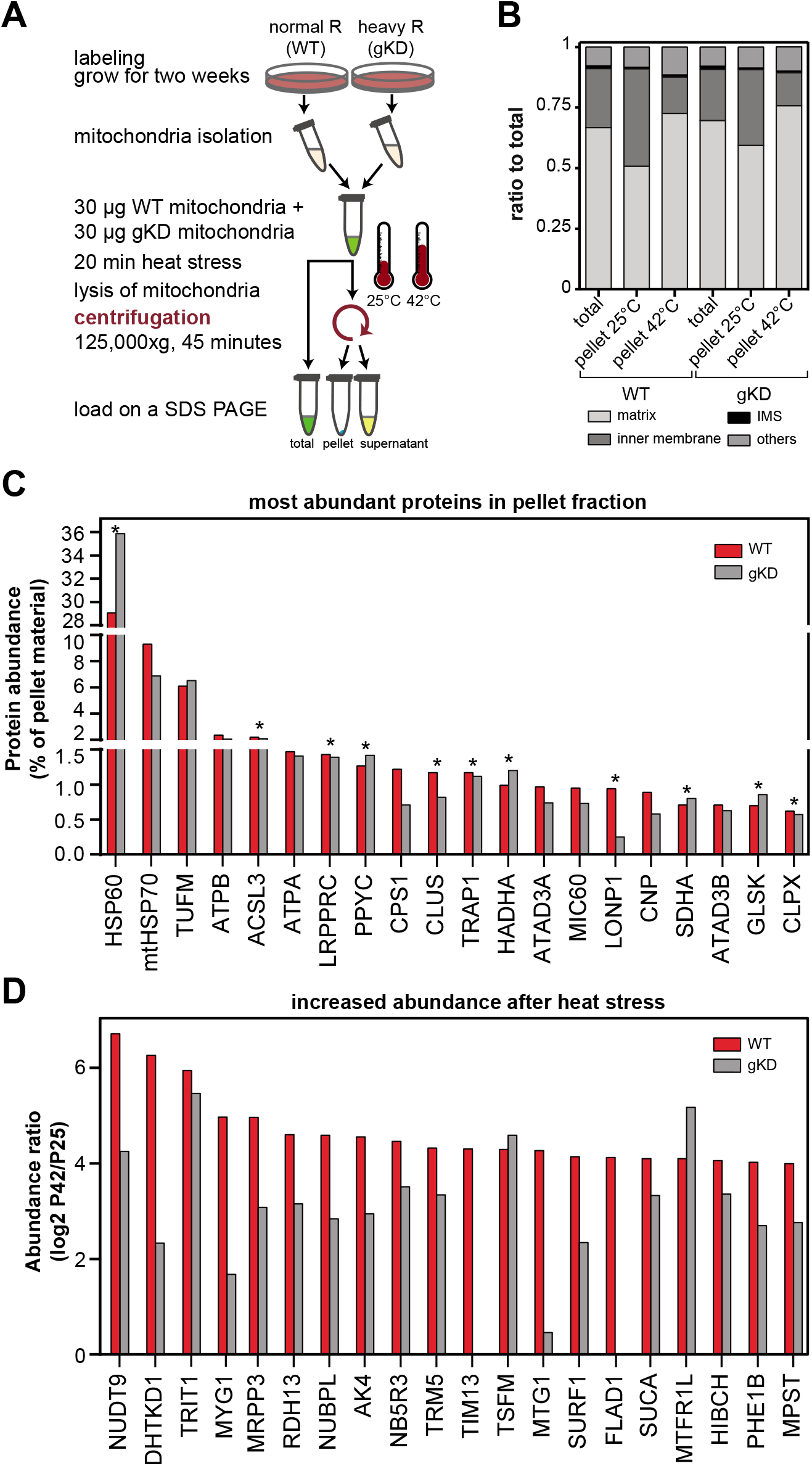
Heat-induced aggregation of mitochondrial proteins. (A) Schematic flow diagram for SILAC labeling and mitochondrial aggregation assay. WT and LONP1 gKD cells were cultured two weeks in medium with normal (WT) or heavy (gKD) isotope-labelled arginine. After isolation, WT and LONP1 gKD mitochondria were mixed 1:1 and treated for 20 min at 25°C or 42°C. Soluble (yellow) and insoluble (blue) proteins were separated via ultra-centrifugation (125,000xg). Unstressed and non-centrifugated sample (total, green) were used for proteome analysis. All samples were separated by SDS-PAGE and analyzed by qMS analysis. (B) Relative distribution of insoluble protein abundances (pellet fraction) in dependence on their sub-mitochondrial localization in matrix (black), inner membrane (dark gray), intermembrane space (IMS; bright gray) or other (gray) from the indicated samples. (C) Shown are the 20 most abundant proteins detected in pellet fraction after 42°C heat stress. Proteins, which were significantly more abundant in pellet fraction after 42°C heat treatment compared to 25°C are marked with stars (representing putative aggregated polypeptides); (WT, red; LONP1 gKD, gray). (D) Shown are the 20 most increased proteins in the heat stress pellet at 42°C compared to 25°C given as log2 values of the abundance ratios 42°C/25°C (WT, red; LONP1 gKD, gray).

In each pellet fraction, we were able to detect about 3200 proteins in total, about 20% of which could be assigned to a mitochondrial localization by UniProtKB database comparison. Again, we restricted our analysis to mitochondrial proteins. In a first general assessment of protein aggregation behavior we noticed a change in abundance of polypeptides in the aggregate pellet correlated to the sub-mitochondrial localization of the identified proteins (Fig. 4B). In the pellet of WT at 25°C, about 51% of total protein abundance was represented by matrix-assigned mitochondrial proteins, while after heat stress with 42°C the percentage of matrix proteins increased to ~79%. Similar to WT, percentages of matrix proteins increased in LONP1 gKD mitochondria from ~59% after 25°C to ~76% after heat stress with 42°C (Fig. 4B). In comparison, inner membrane components were decreased after heat shock in all samples. The percentages of proteins with other localizations did not change significantly. This confirmed the involvement of the protease LONP1 in the general stress-dependent maintenance of protein solubility in the matrix compartment, as expected.

Due to the high sensitivity of the MS-based polypeptide detection, we were able to directly determine which mitochondrial proteins became insoluble after heat stress and therefore represent candidates for aggregation-prone proteins. As protein components of high-molecular weight complexes also sediment to a certain degree under the chosen centrifugation conditions, we first calculated which proteins appeared at least twice as much in the pellet after heat stress (P42), reasoning that these heat-sensitive polypeptides represent aggregated proteins. In total, we were able to detect 248 proteins in WT mitochondria with a larger than two-fold increase in the 42°C pellet. Not surprisingly the most abundant proteins in pellet fraction after 42°C heat stress were the two mitochondrial matrix chaperones HSP60 and mtHSP70 (Fig. 4C). From a group of the 20 most abundant proteins in the pellet fraction, 11 proteins were significantly more abundant in the pellet after 42°C compared to the pellet fraction after heat-treatment with 25°C (Fig. 4C). Due to their heat-dependent sedimentation behavior we suggest these proteins as aggregation candidates. Overall, the number of different polypeptides in the aggregate pellet did not change significantly by the depletion of the LONP1 protease, however, we detected significant quantitative differences in the aggregate proteome of WT and LONP1 gKD mitochondria (see following paragraph). The high proportion of HSP60 in the aggregate pellet (over 25%) is most probably correlated to its chaperone properties. Other members of the mtPQC system were strongly represented in the aggregate pellet too (TRAP1, LONP1, CLPX (ATP-dependent Clp protease ATP-binding subunit clpx-like, mitochondrial), YMEL1) Also, LONP1 itself represented an abundant component in the aggregate pellet. Due to its depletion in the LONP1 gKD mitochondria, its amounts in the aggregate pellet were not significant anymore. We also found major metabolic enzymes of the mitochondrial matrix represented strongly in the aggregate: e. g. PYC (pyruvate carboxylase), and members of the respiratory chain like SDHA, ATPB (ATP synthase subunit beta, mitochondrial) and ATPA (ATP synthase subunit alpha, mitochondrial). An interesting component of the aggregate is the protein LRPPRC (Leucine-rich PPR motif-containing protein, mitochondrial) that is involved in mitochondrial translation by regulating RNA-stability. The ACSL3 enzyme (long-chain-fatty-acid CoA ligase 3), involved in lipid biosynthesis was also a very abundant component of the aggregate pellet. However, its exact mitochondrial localization is unclear, it may potentially be a component of mitochondria-ER contact sites. The previously identified high aggregation-prone translation factor TUFM (elongation factor Tu, mitochondrial) was very abundant in the aggregate pellet although its amounts were not strongly increased after the heat stress treatment. The strongest heat-dependent aggregation was observed for the proteins NUDT9 (ADP-ribose pyrophosphatase, mitochondrial), DHTKD1 (Probable 2-oxoglutarate dehydrogenase E1 component DHKTD1, mitochondrial), TRIT1 (tRNA dimethylallyltransferase), MYG1 (MYG1 exonuclease), MRPP3 (Mitochondrial ribonuclease P catalytic subunit) with an abundance ratio 42°/25°C of more than 30. These high-aggregation polypeptides typically have a relative low abundance in the pellet fraction (Fig. 4D). Interestingly, three out of these five detected proteins were involved in mRNA or tRNA synthesis and therefore part of the mitochondrial protein biosynthesis. The most abundant proteins found in the top group were TSFM (elongation factor Ts, mitochondrial) in WT and LONP1 gKD, PHE1B (pyruvate dehydrogenase E1 component subunit a) in WT; ACAT1 (acetyl-CoA acetyltransferase), ETFA (electron transfer flavoprotein subunit a) in LONP1 gKD (both important for fatty acid beta-oxidation).

We also performed a direct comparison of the sedimentation behavior of protein in WT versus LONP1 gKD mitochondria to determine the quantitative effect of a LONP1 depletion on the stability of the mitochondrial proteome. We plotted the abundance ratios of heat-aggregated proteins (log2 of quotients of abundance values in the pellet fraction after 42°C heat stress and 25°C control sample) obtained from WT and LONP1 gKD together in a dot blot graph (Fig. 5A). This graph represents the amounts of all detected polypeptides in the aggregated pellet in comparison of WT and LONP1 gKD mitochondria. Protein spots in the lower part of the graph (along the X-axis), exhibited a higher aggregation in WT, while the area in the upper part (along the Y-axis) indicates proteins that showed a higher aggregation in LONP1 gKD mitochondria. Theoretically, dots representing proteins with identical aggregation behavior in WT and gKD mitochondria should be located on the diagonal. As the quantitative measurements intrinsically contain minor variabilities, we defined proteins close to the diagonal as non-significant. For example, LONP1 itself showed a high tendency to aggregate after heat stress (Fig. 5A; yellow dot), but as expected no difference in aggregation in WT and gKD was observable. Most proteins with a low aggregation tendency in both LONP1 gKD and WT were membrane proteins or subunits of membrane complexes like the TIM/TOM translocases or respiratory chain complexes. We found 82 proteins whose aggregation was increased more than 2 times in LONP1 gKD compared to WT (Fig. 5A, red dots), which are potential candidates for a LONP1-dependent aggregation. These proteins have no particular aggregation tendency in WT mitochondria, but show a higher protein amount in the aggregate of LONP1 gKD mitochondria. As more abundant proteins may be also represented more in the aggregate pellets, the analysis only of abundance levels as such may not represent how strong a polypeptide is aggregation-prone per se. Hence, we corrected the abundance values of polypeptides in the aggregate pellet with the overall cellular abundance in LONP1 gKD cells and created a factor representing the true aggregation tendency of each protein. This ‘aggregation factor’ was obtained by a division of the polypeptide aggregation ratio in gKD (Table S3) by the LONP1-dependent abundance change in the total proteome (Table S1), and indicated the true dependence of each polypeptide on LONP1 concerning its solubility under heat stress. Of this group of putative proteins with a strong LONP1 dependence, 27 were located in mitochondrial matrix, nine in the inner membrane and nine proteins were located in the outer membrane or in the intermembrane space: Based on their localization the latter proteins most likely can be excluded as candidates for a LONP1-dependent behavior. The six proteins most dependent on LONP1 concerning their aggregation were located in the left upper area of Fig. 5A, showing no aggregation in WT mitochondria after 42°C heat stress but a high aggregation on LONP1 gKD. This is also represented in their aggregation factor: LACTB2 (endoribonuclease), an enzyme important for mitochondrial translation, showed the highest aggregation factor (Fig. 5B), since its total abundance was reduced to 34% in gKD mitochondria and it highly aggregated gKD but not in WT mitochondria. DDAH2 (dimethylarginine dimethylaminohydrolase 2) showed the highest aggregation tendency in gKD and no aggregation in WT mitochondria. As this protein was only slightly increased in LONP1 gKD cells, the overall aggregation factor was lower than LACTB2. The mitochondrial matrix proteins AASS (mitochondrial α-aminoadipic semialdehyde synthase) and CLPP showed still a high aggregation factor despite a lower abundance in gKD mitochondria (Fig. 5B) and are, like LACTB2 and DDAH2, likely LONP1 targets based on this analysis. TIMM29 (mitochondrial import inner membrane translocase subunit) as a constituent of the inner membrane may represent a possible LONP1 target due to its matrix-facing loop (32) that could serve as a LONP1 interaction site. The total abundances of TRUB2 (mRNA pseudo uridine synthase 2), component of mitochondrial translation, and TXNRD2 (thioredoxin reductase 2), a component of the ROS-stress response, showed no change in gKD cells, but both aggregated more in LONP1 gKD mitochondria after heat stress (Fig. 5B; Table 1). The proteins SLC25A18 (mitochondrial glutamate carrier 2) and MSRB2 (mitochondrial methionine-R-sulfoxide reductase B2) also belonged to the group of potential LONP1 targets. COA6 (cytochrome c oxidase assembly factor 6 homologue), is located in the inter membrane space (33) and is therefore unlikely to be affected by LONP1. One of the proteins with a low aggregation factor in our experiment is QRSL1 (mitochondrial glutamyl-tRNA aminotransferase subunit A). It showed a high aggregation tendency after temperature stress, but while it was increased also more than eight times in LONP1 gKD mitochondria under steady-state conditions, it had one of the lowest aggregation factors (Fig. 5B).

**Figure 5.**
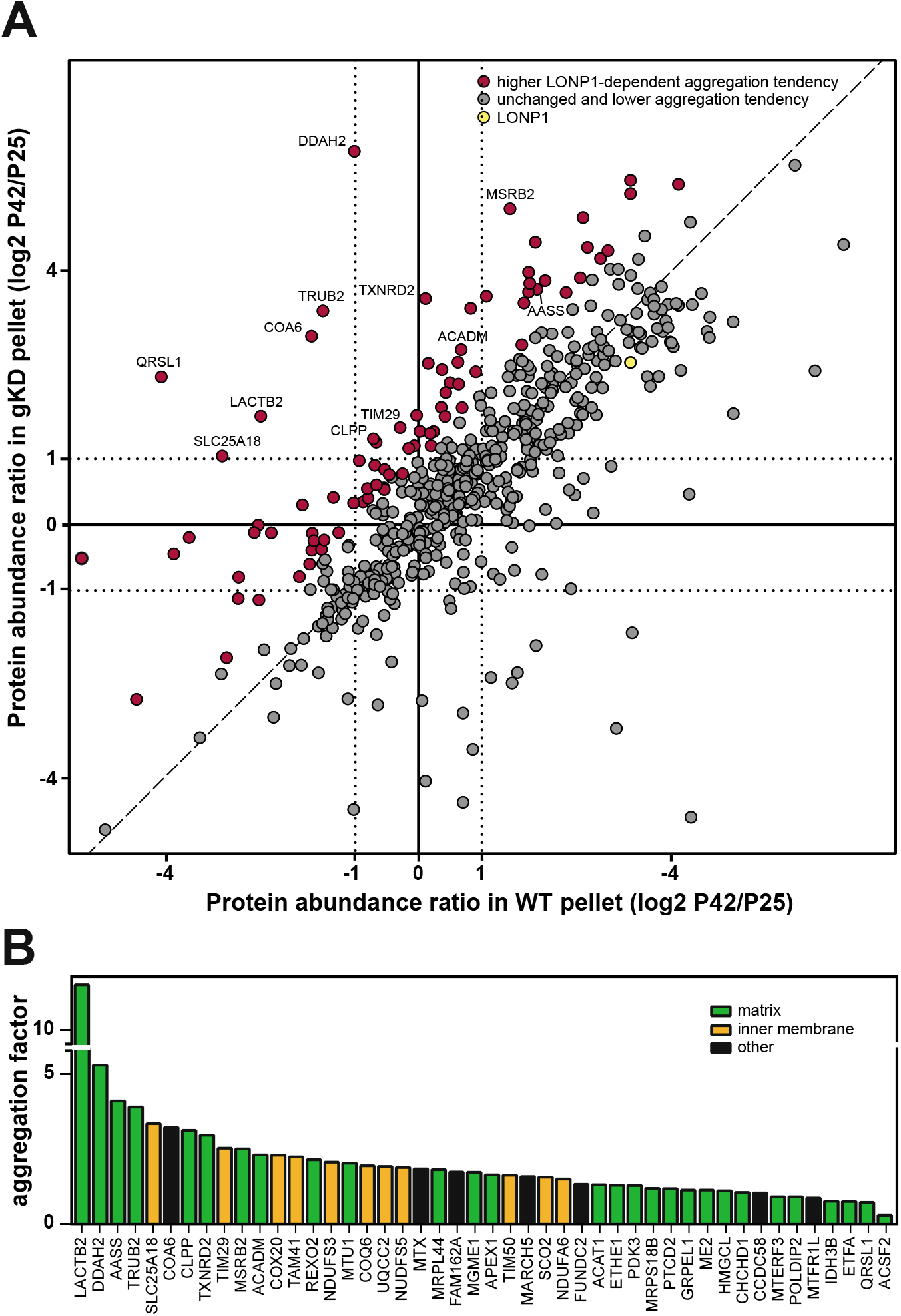
Effect of LONP1 gKD on the heat-induced aggregation in mitochondria. (A) Comparison of the aggregation behavior of mitochondrial proteins in LONP1 gKD and WT on the proteome level. Shown are the log2 values of the abundance ratios of proteins in the high-speed centrifugation pellets at 42°C divided by that at 25°C obtained by qMS from WT and LONP1 gKD mitochondria. Proteins with increased aggregation in WT are represented on the right half of the graph (X-values >1) while proteins with increased aggregation in LONP1 gKD are found in the upper part of the graph (Y-values >1). Proteins with a significant LONP1-dependent aggregation behavior are labeled red (aggregation ratio gKD/WT >2), other proteins are shown in gray. Spots near the dashed line (slope of 1) represent proteins with identical aggregation behavior (ratio P42°/P25°) in WT and LONP1 gKD. LONP1 is shown as yellow dot. (B) Proteins with highest aggregation factor, indicating the LONP1-dependence of their aggregation normalized to the overall abundance. Only proteins are considered that show significant aggregation (ratio P42°/P25° >2) in gKD mitochondria. Matrix proteins (green), inner membrane proteins (orange), proteins in other localization (black).

In order to confirm the qMS results, we performed the aggregation assay with a classical Western blot detection using available different antibodies (Fig. S2). As expected, the matrix proteins LONP1, ACO2, HSP60 and TSFM showed no aggregation behavior in unstressed samples. After heat stress at 42°C, LONP1, HSP60 and TSFM were detected mostly in pellet fraction. ACO2 did not aggregate completely after heat stress at 42°C, but after 45°C almost ~90% ACO2 was found in pellet fraction. After 37°C incubation, TUFM was mostly detectable in pellet fraction. The membrane protein TIMM23 was used as solubility control, it remained in supernatant fraction under normal as well as heat stress condition (Fig. S2). As the Western blot analysis was restricted to relatively abundant reporter proteins, no clear changes in aggregation behavior between WT and LONP1 gKD were observable, similar to the qMS data.

Taken together, we have identified a group of novel aggregation-prone proteins whose accumulation as misfolded species under heat stress is prevented LONP1. These proteins perform important roles for normal mitochondrial functionality, in particular for the expression of mitochondrial-encoded proteins. In contrast to the more abundant components of the PQC system, in particular the chaperones of the HSP60 and HSP70 protein families, LONP1 did not seem to play a major role in maintaining the solubility of highly abundant metabolic enzymes in mitochondria.

## Discussion

Mitochondrial dysfunction has been recognized as a major factor in many human pathologies, in particular, due to its accumulative fashion, in neurodegenerative diseases (34,35). Moreover, protein misfolding and subsequent aggregation is a main cause of functional defects in mitochondria and is counteracted by the activity of the mtPQC machinery (36). ATP-dependent protease enzymes in the matrix compartment, in human mitochondria represented by the protein LONP1, are prominent members of this mtPQC system (37).

### Effects of reduced LONP1 protein levels

A complete genetic knock out of the LONP1 protease was not viable in cultured cells (23,38) and resulted in embryonic lethality in mice (25). Therefore, we chose to create a CRISPR/Cas9 mediated genetic knock down of LONP1. The LONP1 genomic knock down cells showed a stable reduction of LONP1 protein level to about a quarter of the WT levels. To analyze the adaptive effects of this LONP1 reduction, we investigated morphology and diverse functions of mitochondria isolated from the novel cell line. In general, the cells showed no observable morphological phenotype with essentially intact mitochondria cristae structures and a healthy tubular mitochondria network. This was in contrast to a previous publication that used a siRNA-mediated knock down of LONP1 showed a fragmented mitochondrial network (38). However, a doxycycline-induced expression of the LONP1 siRNA was used in this study. As doxycycline is also a mitochondrial toxin (39) it is unclear if the observed defects were directly connected to the reduction of LONP1 amounts or caused by a general functional defect due to the toxin treatment. Furthermore, the protein amount of SOD2, a ROS-stress response protein, increases after doxycycline treatment, indicating that doxycycline generates oxidative stress in mitochondria (39). Since LONP1 expression is also up-regulated by ROS stress (7,22), the use of doxycycline as an inducer of a LONP1 knock down seems problematic. Other studies with a very strong depletion of LONP1, either after a siRNA-mediated LONP1 knock down in human fibroblasts (23) with loss of more than ~95% of LONP1 as well as a complete deletion of the protease homolog Pim1 in yeast (14) exhibited a stronger phenotype, e. g. concomitant a complete loss of mitochondrial cristae structure. A complete loss of LONP1 function has a very negative effect on cell survival as has been observed by the embryonal lethality in LONP1 knock out mice and the higher apoptosis rate in cells with almost complete down-regulation of LONP1 (23,38). However, as our LONP1 gKD cells does not rely on the use of external mitochondrial toxins and a milder depletion of the protease, the model is more suitable to study functional effects that occur in pathologies related to a reduced, but not completely abolished LONP1 function.

By observing a reduction of the protein amount of LONP1, we would also expect a reduction of the degradation rate to a similar percentage. However, with imported reporter proteins we were able to detect only a subtle decrease in degradation rates. This suggests that in WT mitochondria not all LONP1 complexes are occupied with substrate degradation or that they are able to degrade the reporter proteins more quickly. While in LONP1 gKD mitochondria the majority of LONP1 proteases are occupied, it takes significantly more time to degrade the reporter proteins. Our analysis therefore shows that the reduced LONP1 amount is sufficient to ensure a basal degradation efficiency under normal conditions, where not many misfolded polypeptides accumulate. If the mitochondria, both WT and LONP1, are now treated with menadione, there are more proteins damaged by free oxygen radicals and thus more LONP1 substrates are available. Hence, under stress conditions hardly any reporter proteins were degraded. In addition, a slightly reduced degradation rate in LONP1 mitochondria could be related to the decrease in mitochondrial ATP concentration in LONP1 gKD mitochondria, as the AAA+ protease requires the hydrolysis of ATP for its function. The reduction of mitochondrial ATP concentration in the LONP1 gKD cells is likely due to a defect in oxidative phosphorylation rates, possibly correlated to the reduced translation of the mitochondrial-encoded ATPase subunit ATP8, which we observed here. In addition, a LONP1 siRNA knock down study also showed a significant reduction in respiration in fibroblasts (23) and a similar decrease in the ATP concentration was observed in heterozygous LONP1 knock out mice (25,40). Also in yeast, the depletion of Pim1 shows a strong impairment of cellular respiration and a complete loss of membrane potential (14).

In our study LONP1 gKD mitochondria showed no increased ROS accumulation, suggesting that under non-stressed conditions the residual LONP1 amounts were enough to maintain mitochondrial functions, in particular for the oxidative metabolism. However, LONP1 gKD cells reacted more sensitive to external ROS stress in comparison to WT cells. It is likely, that the reduced amounts of the protease are unable to cope with increased amounts of oxidatively modified polypeptides that occur under stress conditions, eventually leading to a failure of mitochondrial functions. This correlates well with previous observations in yeast, where a *PIM1* deletion mutant was also more sensitive to ROS treatment (10). Interestingly, levels of LONP1 and SOD2, the main ROS scavenger enzyme in mitochondrial matrix, seem to be correlated, since in our LONP1 gKD cells SOD2 levels were decreased, and vice versa SOD2 heterozygote knock out mice showed a significantly lower LONP1 level, especially at an older age (41).

Interestingly, the other mitochondrial matrix protease CLPP was found to be reduced in LONP1 gKD cells. This seemed to be counterintuitive since it would be expected that a lack of LONP1 may need to be compensated by an increased amount of CLPP. Indeed, analysis of a doxycycline-dependent LONP1 siRNA knock down showed, not surprisingly, an increase CLPP protein level (42). In the light of a reduced level of LONP1, a concomitant reduction of the potentially alternative protease CLPP would certainly represent a disadvantage for the cell, as CLPP and LONP1 share some substrate proteins, including oxidative phosphorylation, TCA cycle, amino acid and lipid metabolism (43). In a recent study in human cancer cells, it was observed that the expression of LONP1 and CLPP was strongly related. The genes for both matrix proteases are located close to each other on chromosome 19 and are co-expressed (43). Hence, it is possible that the deletion in the promoter region of LONP1 in our gKD cells also lead to a poorer expression of CLPP.

While analyzing the changes in the mitochondrial proteome after LONP1 gKD, we found a few proteins, which were significantly up- or down-regulated. This likely indicates some degree of proteome remodeling as a potential adaption mechanism to the reduction of the protease levels. The abundance of common LONP1 substrates, like ACO2 (21) and TFAM (44), was not changed, another MS study in heterozygote LONP1 knock out mice (45) showed no changes in TFAM or ACO2 levels either. The lack of effect suggests that reduced LONP1 protein levels are sufficient to degrade ACO2 and TFAM under normal conditions. Although not yet considered as potential substrate proteins, specific members of the preprotein import-related HSP70 chaperone system (HSCB, DNAJC30, GRPEL2) and also components the membrane-integrated TIM/TOM translocase complexes were strongly increased after the LONP1 gKD, although no obvious import defect was detectable in LONP1 gKD mitochondria. As it is unlikely that in particular the TIM/TOM components are genuine LONP1 substrates, the up-regulation of HSP70 co-chaperones GRPEL2 and DNAJC19 might indicate the possibility of adaptation processes in the gKD cells in order to normalize import rates of mitochondrial proteins in case of a lack of the protective LONP1 protease. The increase in TIM/TOM receptors components TIMM21 and TOMM20 might also help to recognize and interact better with presequence-containing mitochondrial proteins and thereby also increase import efficiency. At least in the case of the mtHSP70 co-chaperones it is likely that a reduction of LONP1 might cause a mild accumulation of misfolded polypeptides even under normal conditions and that the increased amount of co-chaperones helps the mtHSP70 to compensate this problem.

The components of the intramitochondrial protein translation system also seemed to be slightly altered in LONP1 gKD cells. These minor effects may be caused by an adaptive behavior of the gKD cells and a result of decreased amounts of certain translation-related components. Two of them, GATB and PARS2, are involved in the synthesis of aminoacyl-tRNAs and another, MRPS11, is a structural protein of the mitochondrial ribosome. It is unclear why these specific proteins have reduced amounts in response to a LONP1 reduction although their decrease might explain the observed reduction of the translation of certain mitochondrially-encoded proteins in LONP1 gKD cells.

In summary, the LONP1 gKD cells show a subtle proteome remodeling, both reductions and increases in specific mitochondrial proteins that likely reflects and adaptation to the partial loss of LONP1-related PQC properties in these mitochondria. It seems unlikely that an increased level of specific proteins simply indicates a reduced degradation, at least under normal growth conditions.

### LONP1-dependent aggregation behavior

As LONP1 has been described as a major component of the mitochondrial protein homeostasis machinery in particularly under stress conditions, we aimed at to confirm this claim by a comprehensive proteome-wide characterization of protein aggregation in mitochondria. In a previously published analysis heat-dependent aggregation reactions in mitochondria isolated from HeLa cells (6), based on 2D electrophoresis, we have already identified a few highly aggregation-prone polypeptides. However, for technical reasons, the previous approach focused on the soluble proteome before and after heat stress treatment. In contrast, the mass-spectroscopic approach used here allowed for the first time a direct identification and also quantification of proteins from the aggregate pellet. In addition, the combination of aggregate characterization with the analysis of proteome changes in the LONP1 gKD cells yielded direct and detailed insights into the role of this protease during heat stress conditions.

In our previous study we observed, with few remarkable exceptions, that the soluble mitochondrial proteome did not exhibit drastic quantitative changes in response to heat-induced stress conditions (6). In contrast, the qMS-based analysis of the aggregate pellet in this study resulted in a high number of proteins with a considerable aggregation tendency. In addition, a decrease of the mtPQC protease LONP1 exacerbated the aggregation rates of many proteins. As the LONP1 protease is able to degrade misfolded proteins in the matrix compartment (46,47), this confirms directly on a proteomic level that LONP1 fulfills an aggregation-protective role in mitochondria under heat stress. It is therefore highly conceivable that the excess aggregated polypeptides represent potential substrates of the LONP1 protease, providing additional information to determine its substrate range.

Many of the most abundant proteins in the aggregation pellet belonged also to the proteins with the highest overall abundance in mitochondria – represented by the total mitochondrial protein amount. This indicates that an aggregation reaction becomes more likely the higher the amount of misfolded polypeptides is in a certain cellular microenvironment. These very abundant proteins end up in the aggregate pellet largely independent of the activity of LONP1 as the total amount of LONP1 complexes is too low to cope with the high number of potentially misfolded polypeptides derived from these proteins under stress conditions. In contrast, we also observed a strong aggregation tendency for some proteins with a lower overall abundance. In this case, their aggregation behavior often was considerably influenced by the available amounts of the LONP1 protease. This group of proteins likely represented intrinsically thermo-labile polypeptides that are the most likely targets of the LONP1-mediated degradation.

Interestingly, classical chaperone proteins like HSP60, mtHSP70, CLPX were major components in the aggregate pellet. This also correlates with their overall high total abundance inside mitochondria. Their presence in aggregate pellet is likely based on their typical binding affinity to misfolded proteins, indicating that they were pulled down together with the majority of destabilized polypeptides. This also reveals that chaperone binding as such is not necessarily successful in keeping a substrate polypeptide in a soluble state during heat stress conditions. It is possible that CLPX is not only active as a chaperone subunit of CLPXP protease complex (16) within the matrix but also has a possible chaperone function on its own, interacting with misfolded proteins and thus potentially co-aggregating. This is supported by our observations that indicate a roughly 5-7 times higher total amount of CLPX than CLPP in mitochondria. Interestingly, yeast mitochondria have lost the CLPP protease enzyme but still retain the corresponding chaperone component Mcx1 (48). Similar to the chaperone enzymes, also LONP1 itself was relatively strongly represented in the aggregate pellet. Although LONP1 may have some chaperone properties itself (49), it seems to be a heat-labile protein that strongly aggregates and is 4-5 times more abundant in the pellet after heat stress, both in WT and the gKD cells (irrespective of the lower concentration in the mutant mitochondria). This rather strong aggregation of LONP1 had already been observed in our earlier study (6). Concerning the presence of chaperone enzymes in the aggregate pellet, it is important to note that the chaperones themselves may exhibit a certain instability. In particular in case of large oligomeric protein complexes, like HSP60 and also LONP1, as they may dissociate and the polypeptides in monomeric form may become unstable and aggregate under stress. Indeed, HSP60 in yeast has been already identified as one of the major degradation substrates of the Pim1 protease at elevated temperatures (50).

The translation factor TUFM represented a major component of the aggregate pellet, confirming previous observations of our lab, that its amounts are depleted strongly from the soluble proteome after heat stress (6). Indeed, its partner protein TSFM, the associated GTP exchange factor, belonged to the proteins with the highest aggregation tendency (abundance ratio 42°C/25°C) in mitochondria. We had previously proposed that an enhanced aggregation-sensitivity of the translation factor serves as a stress-reactive switch that shuts down mitochondrial protein biosynthesis in reaction to heat shock (6). This should reduce the risk of newly synthesized nascent mitochondrial proteins contributing to aggregate formation while the mitochondria are experiencing and recovering from heat shock.

We have also identified ACSL3 (Long-chain-fatty-acid--CoA ligase 3), as a protein that is one of the largest contributors to the pellet after heat stress. Although ACSL3 is probably localized to the outer mitochondrial membrane, the larger part of the protein is facing the cytosol and participates in lipid biosynthesis (51). ACSL3 (and its relative ACSL4) possibly is a structurally unstable protein that is highly aggregation-prone under elevated temperatures. Correlating with its anchoring in the outer membrane, aggregation behavior of ACSL3 is likely independent of the function of LONP1.

The direct dependence of the aggregation behavior of individual polypeptides was expressed in form of an “aggregation factor” that also takes the total abundance changes in the LONP1 gKD cells into account as some proteins with high aggregation tendencies were also more abundant in gKD cells. Most proteins with a higher aggregation factor fall into the functional groups of RNA maturation, β-oxidation, or protein assembly enzymes. One of the most prominent examples is LACTB2, which showed a very high LONP1-dependent aggregation tendency after heat stress. LACTB2 is involved in mtRNA processing and essential for transcription in mitochondria. A siRNA-mediated reduction of LACTB2 showed an accumulation of mitochondrial transcripts (52). As a reduction of LACTB2 levels exhibited severe cellular defects, it is possible that the accumulation of mRNA within the matrix may have a negative influence on the translation and eventually on mitochondrial function per se. Therefore, the reduction of LACTB2 in LONP1 gKD cells may also explain a decrease in mitochondrial translation rate, as is observed in this study. The strong dependency of LACTB2 on LONP1 function fits to previous observations that the protease is either directly or indirectly involved in mitochondrial gene expression, as it may act as transcription regulator via TFAM (44) and binds to mtDNA in promoter regions (53). A faster LONP1-dependent aggregation of LACTB2 and TRUB2 could also contribute to the stress-protective shutdown of mitochondrial protein expression in addition to the inactivation of TUFM/TSFM, as mentioned above.

It has been shown, that a loss of LONP1 leads to an accumulation of unprocessed forms of a few imported proteins e. g. CLPX in the mitochondrial matrix, potentially due to an inhibition or inactivation of the mitochondrial processing protease (26). These pre-proteins may be structurally instable and aggregate faster with a higher tendency in LONP1 gKD mitochondria already under normal conditions. Both in the yeast *pim1*Δ deletion strain (14) as well as in a study using a siRNA-mediated reduction of the LONP1 protease (23) an accumulation of electron-dense particles - putatively protein aggregates - have been observed already under normal conditions. However, in case of CLPX, our observations indicated that CLPX exhibits a high aggregation tendency in general, which seems independent LONP1.

A puzzling finding is the high aggregation propensity of COA6 (cytochrome c oxidase assembly factor 6 homologue) specifically in the LONP1 gKD mitochondria as this protein is normally localized in the intermembrane space (33). This shows a clear LONP1 dependence. One possible explanation for this outlier is that COA6 may be localized in the matrix during the biogenesis pathway and/or mis-localized into the matrix in LONP1 gKD cells. Thus, COA6 cannot be excluded as a LONP1 substrate but needs a more detailed future examination.

Interestingly, we found some membrane proteins with a high LONP1-dependent aggregation factor. As a protease localized in the matrix compartment it is so far unclear how much LONP1 would contribute to the degradation of membrane proteins of the inner membrane. However, it cannot be excluded that certain membrane proteins could be possible substrates of LONP1, as they contain a matrix-exposed domain, serving as a degradation or at least a chaperone target. The TIM translocase subunit TIMM29 showed a high aggregation tendency and its amount reduced in LONP1 gKD mitochondria. As the membrane protein TIMM29 contains a prominent matrix loop (54) it may indeed be a suitable target for the matrix protease LONP1. Although integral membrane proteins as such are mostly protected from aggregation there are also membrane proteins that have a very large non-membrane domain, which in turn may make them vulnerable to aggregate formation. For example, ACSL3 has only a small anchorage segment in the outer mitochondrial membrane (51), but the majority of the protein is exposed to a water environment, possibly causing the strong aggregation propensity.

It has been suggested that larger proteins have higher aggregation tendencies due to their more complicated structural complexity. However, we could not find a correlation between molecular weight and aggregation tendency. E. g. a very small protein with 15 kDa (COA6) and as well as larger proteins like QRSL1 with 57 kDa both showed a high aggregation tendency.

### LONP1 gKD cells as aging model

Mitochondria are the major cellular source of ROS and the generation of free radicals increases with age (55,56) thereby contributing to the decline of cellular functions. Elevated ROS concentrations typically lead to oxidation of amino acid side chains in polypeptides, subsequently leading to defects in protein structure, misfolding and potentially aggregation (57,58). LONP1 is the only known protease which degrades oxidatively modified proteins (21) in the mitochondrial matrix and thereby prevents the accumulation of these damaged polypeptides. However, activity (59), mRNA level (60) and protein amounts of LONP1 decreases while aging in most tissues (61). In old murine muscle cells, LONP1 protein levels were reduced about fourfold (41), which correlates well to the LONP1 protein level in our gKD model. In aging rat liver cells LONP1 has decreased proteolytic activity, which leads to an accumulation of damaged proteins, although the protein level of LONP1 itself did not change (59). Correlating with this effect, mitochondrial functionality is reduced, representing a major factor in the emergence of neuronal diseases (62–64). With their relatively mild phenotype, the LONP1 gKD cells retain basic mitochondrial functionality but exhibited a higher sensitivity to ROS stress and a lower ATP content, typical hallmarks of aging cells (56). In contrast to conventional transient siRNA-knock down experiments, our cell model has adapted to a reduction of LONP1 and better reflects the long-term alterations taking place in aging cells. In summary, our newly designed LONP1 gKD cell model essentially replicates the above-mentioned behavior of the protease in aging cells and can therefore be potentially used as an experimental model for age-related pathological processes.

## Supporting information

Supplemental Table 1 Protein total abundances

Supplemental Table 2 Protein aggreggate abundances

Supplemental Table 3 Protein aggregation factors

## Acknowledgements

We would like to thank especially M. Fuhrmann and B. Gehring for their technical support of the project. Dr. A. Heimbach from the Next Generation Sequencing Core Facility of University Hospital Bonn performed the genomic sequencing of the LONP1 gKD cells. In addition, we would like to thank C. Runz, L. Lüdecke, L. Ruland, D. Putcha-Schomberg, Drs. G. Cenini, and W. Jaworek for their critical and constructive discussion of the manuscript.

## Abbreviations

AAA+: ATPases associated with a wide variety of cellular activities
Δψ: mitochondrial membrane potential
gKD: genetic knockdown
HSP: heat shock protein
m: mature form
mt: mitochondrial
p: precursor form
PQC: protein quality control
qMS: quantitative mass spectrometry
ROS: reactive oxygen species
SILAC: stable isotope labeling with amino acids in cell culture
siRNA: small interfering RNA
TIM: preprotein translocase complex of the inner membrane
TMRE: tetramethylrhodamine
TOM: preprotein translocase complex of the outer membrane
UPR^mt^: mitochondrial unfolded protein response
WT: wild type.

