## Supplemental figures for "Proteomic analysis of the role of the quality control protease LONP1 in mitochondrial protein aggregation"

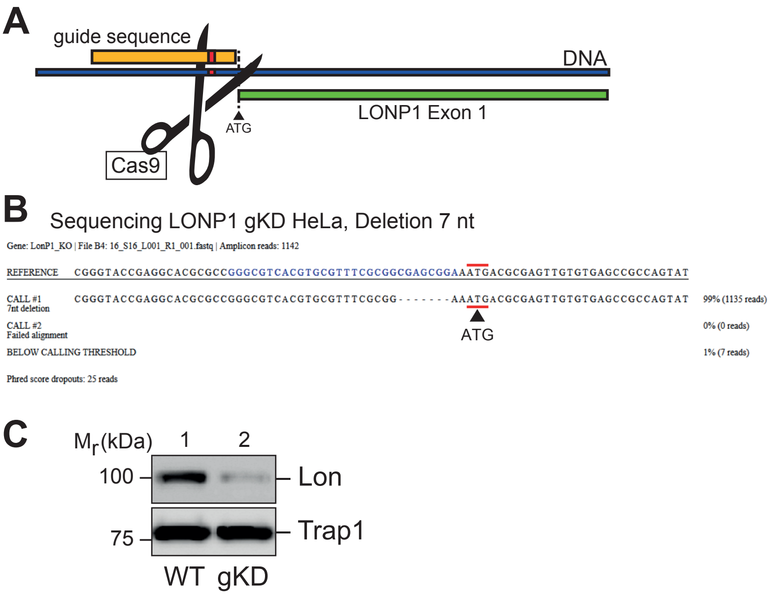


**Figure S1 CRISPR/Cas-mediated Lon knockdown and sequencing**

(A) Schematic diagram of the CRISPR/CAS9-mediated genetic knockdown procedure, Binding site of the guide sequence (orange) upstream the LONP1 start codon (green) and interface of the endonuclease CAS9 (red). (B) Sequence data of the used LONP1 gKD clone obtained by MiSeq-sequencing. The used guide sequence is marked in blue and the start code ATG of LONP1 is labeled red. (C) Verification of the reduction of LONP1 protein level in HeLa wild type (WT) and LONP1 gKD mitochondria analyzed by SDS-PAGE, Western blot, and immunodetection using the indicated antibodies. Trap1 signals were used as loading control.

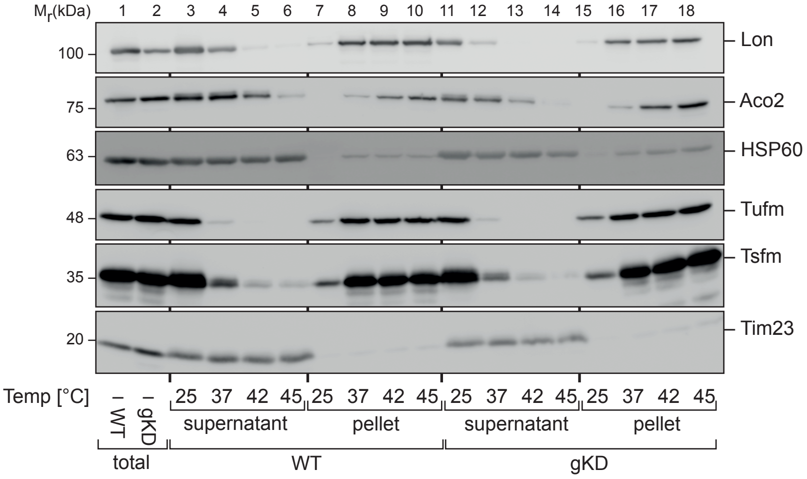


**Figure S2 Heat-induced aggregation of mitochondrial proteins**
Analysis of protein aggregation behavior in isolated mitochondria after 20 min heat stress of mitochondria at different temperatures. Aggregated proteins were sedimented from the detergent-lysed mitochondria by ultra-centrifugation at 125,000xg and detectable in the pellet fraction, soluble proteins remain in the supernatant. Total cell lysates (total) were not stressed. All samples were analyzed by SDS-PAGE, Western blot and immunodetection with the indicated antibodies. Here, the signal of TUFM was used as aggregation control and TIM23 as control for soluble proteins.
